# Volatilomic complexity of three Northern Greenland bacterial isolates across a salt gradient

**DOI:** 10.1101/2025.06.23.660987

**Authors:** Miguel Ángel Salinas-García, Kajsa Roslund, Mathias Bygum Risom, Anders Priemé, Riikka Rinnan

## Abstract

The High Arctic deserts of remote northern Greenland are expected to become warmer and wetter due to climate change. Precipitation changes will increase fluctuations in surface soil salinity, and the same happens for thawed permafrost soil where stable salt concentrations are replaced with fluctuating salinity during annual freeze-thaw cycles. Both have unknown effects on the microbial communities and their emissions of microbial volatile organic compounds (MVOCs). These compounds are produced from various pathways mainly as secondary metabolites and have ecological and climatic implications when released into the environment and the atmosphere. Thus, it is important to explore the effects of environmental changes, such as changes in salinity, on soil microbial communities and their MVOC emissions. Here, we characterize the MVOC production of three novel bacterial isolates from northern Greenland throughout their growth period under low, moderate, and high salt concentrations. We demonstrate that salinity significantly alters both the quantity and composition of MVOCs emitted by all three strains, including changes in the emissions of sulphur- and nitrogen-containing compounds, potentially leading to ecosystem nutrient loss. The observed changes in MVOC profiles suggest that changes in soil salinity due to climate change could alter microbial metabolism and MVOC emissions, with potential implications for Arctic nutrient cycling and atmospheric chemistry.

## INTRODUCTION

Microbes produce volatile organic compounds (MVOCs) for various purposes, such as communication and competition^1^, or as metabolic byproducts (e.g., glucose fermentation into ethanol). VOCs play a role in atmospheric processes, as they can oxidise in the atmosphere to form secondary organic aerosols or increase tropospheric ozone, indirectly increasing the lifetime of methane^2^. MVOCs can be particularly important in terrestrial ecosystems with reduced plant cover, such as the Greenlandic Ice Sheet^3^ and barren permafrost regions^4^, where MVOC emissions are largely unexplored and dependent on the composition of their microbial communities.

Salinity is known to influence the production of MVOCs, such as methanethiol, in estuarine settings^5^, and snowmelt has been shown to induce the production of MVOCs, e.g. methanol, in alpine soils^6^. However, no studies to date have investigated the MVOC production of Arctic soil bacteria under different salinity. In this study, we examined the effects of salinity on the volatilomes (the collection of volatile compounds) of three bacterial strains isolated from different environments of Peary Land. Peary Land in northern Greenland is a polar desert characterised by low precipitation and temperatures^7^. The region is mostly barren, with sparse vegetation in certain areas, and soils are generally poor, saline, and alkaline^8^. However, like the rest of the Arctic region, Peary Land is expected to warm faster than the global average^9^, and the frequency of extreme weather conditions, such as periods of heavy precipitation or drought, may increase^10^. Consequently, these environmental changes could lead to fluctuations in soil salinity and moisture, potentially affecting the microbial communities in these extreme environments. As microbial activity is closely linked to environmental factors, changes in temperature and salinity likely alter metabolic processes, including the production of MVOCs. Understanding how Arctic bacteria adapt to these changes is crucial for predicting potential impacts on nutrient cycling and future MVOC emissions in this region.

The main aim of this study was to characterize and quantify MVOC emissions from the three bacterial strains, using a combination of proton-transfer-reaction time-of-flight mass spectrometry (PTR-ToF-MS) and gas chromatography-mass spectrometry (GC-MS), for both real-time monitoring and identification of a wide range of MVOCs. The strains studied were *Nesterenkonia aurantiaca* CMS1.6, *Oceanobacillus sp.* CF4.6, and *Arthrobacter sp.* KK5.5, isolated from Cape Morris Jessup biological soil crust, Citronen Fjord biological soil crust, and Kap København permafrost, respectively. By analysing their volatilomes across different salt concentrations, we aimed to test two key hypotheses: (1) salinity affects the volatilomes produced by the three Arctic bacterial strains beyond lowered emissions expected from reduced growth alone; and (2) the physiological and metabolic processes underlying MVOC production are affected differently by salinity among the three strains. This study contributes to a broader understanding of microbial adaptations in extreme environments and may have implications for nutrient cycling and volatile emissions in a changing Arctic climate.

## RESULTS

### Identification of MVOCs

We discovered a total of 104 mass-to-charge (*m/z*) signals in the real-time PTR-ToF-MS measurements, corresponding to VOCs produced by the studied bacteria and the nutrient media. **Table 1** lists the tentative identities of the 23 most abundant signals produced by the bacteria that differ from those in the control samples (pure nutrient media). The signals were classified according to their molecular formula: “Hydrocarbon” (compounds containing C and H), “oxygen-containing” (C, H, and O), “sulphur-containing” (C, H, and S), “nitrogen-containing” (C, H, and N), “halogen-containing” (C, H, and a halogen) or “other” (C, H, and two or more different heteroatoms). Further confirmation of compound identities was achieved through GC-MS analysis.

**Table 1.** Most abundant volatile signals produced by the three studied Arctic bacteria. Tentative identifications are based on the observed masses of the PTR-ToF-MS signals, the results from GC-MS measurements, and available literature^11,12^ . Fragment ions are expressed in grey italics. *It should be noted that the signals classified as “hydrocarbon” were not necessarily indicative of the presence of hydrocarbon parent compound, as other heteroatom-containing compounds can break into “hydrocarbon” fragments during the ionisation process.

| SIGNAL (U) | MOLECULAR FORMULA | TENTATIVE IDENTIFICATION | THEORETICAL MASS (U) | MASS DIFFERENCE (U) | IDENTIFIED WITH GC-MS OR IN GAS STANDARD | CLASS |
| --- | --- | --- | --- | --- | --- | --- |
| 31.0178 | (CH<sub>2</sub>O)H<sup>+</sup> | Formaldehyde | 31.0183 | 0.0005 |  | Oxygen-containing |
| 33.0335 | (CH<sub>4</sub>O)H<sup>+</sup> | Methanol | 33.0340 | 0.0005 | X | Oxygen-containing |
| 34.9950 | (H<sub>2</sub>S)H<sup>+</sup> | Hydrogen sulphide | 34.9955 | 0.0005 |  | Sulphur-containing |
| 41.0386 | (C<sub>3</sub>H<sub>5</sub>)<sup>+</sup> | *Common alkenyl fragment, e.g., from isoprene* | 41.0391 | 0.0005 |  | *Hydrocarbon\** |
| 42.0338 | (C<sub>2</sub>H<sub>3</sub>N)H<sup>+</sup> | Acetonitrile | 42.0343 | 0.0005 | X | Nitrogen-containing |
| 43.0542 | (C<sub>3</sub>H<sub>7</sub>)<sup>+</sup> | *Common alkyl fragment, e.g., from 2- and 3-methylbutanol* | 43.0547 | 0.0005 |  | *Hydrocarbon\** |
| 45.0330 | (C<sub>2</sub>H<sub>4</sub>O)H<sup>+</sup> | Acetaldehyde | 45.0340 | 0.0010 | X | Oxygen-containing |
| 47.0128 | (CH<sub>2</sub>O<sub>2</sub>)H<sup>+</sup> | Formic acid | 47.0133 | 0.0005 |  | Oxygen-containing |
| 47.0491 | (C<sub>2</sub>H<sub>6</sub>O)H<sup>+</sup> | Ethanol | 47.0497 | 0.0006 |  | Oxygen-containing |
| 49.0107 | (CH<sub>4</sub>S)H<sup>+</sup> | Methanethiol, with contribution from DMDS and DMTS. | 49.0111 | 0.0004 |  | Sulphur-containing |
| 57.0699 | (C<sub>4</sub>H<sub>9</sub>)<sup>+</sup> | *Common alkyl fragment, e.g., from 1-butanol and 2-methyl-1-propanol* | 57.0704 | 0.0005 |  | *Hydrocarbon\** |
| 59.0491 | (C<sub>3</sub>H<sub>6</sub>O)H<sup>+</sup> | Acetone | 59.0496 | 0.0005 | X | Oxygen-containing |
| 69.0699 | (C<sub>5</sub>H<sub>8</sub>)H<sup>+</sup> | Isoprene, with contribution from 2- and 3-methylbutanal fragments | 69.0704 | 0.0005 | X | Hydrocarbon* |
| 71.0491 | (C<sub>4</sub>H<sub>7</sub>O)<sup>+</sup> | *Common carbonyl fragment, e.g., from 3-methylbutanal* | 71.0496 | 0.0005 |  | *Oxygen-containing* |
| 71.0820 | (C<sub>5</sub>H<sub>10</sub>)H<sup>+</sup> | *Fragment of 2- and 3-methylbutanol and other 5C alcohols* | 71.0860 | 0.0040 | X | *Hydrocarbon\** |
| 73.0648 | (C<sub>4</sub>H<sub>8</sub>O)H<sup>+</sup> | 2-butanone and butanal | 73.0653 | 0.0005 | X | Oxygen-containing |
| 81.0699 | (C<sub>6</sub>H<sub>8</sub>)H<sup>+</sup> | *Fragment of monoterpenes and larger terpenoids* | 81.0700 | 0.0001 |  | *Hydrocarbon\** |
| 87.0804 | (C<sub>5</sub>H<sub>10</sub>O)H<sup>+</sup> | 2- and 3-methylbutanal | 87.0809 | 0.0005 | X | Oxygen-containing |
| **89.0597** | (C<sub>4</sub>H<sub>8</sub>O<sub>2</sub>)H<sup>+</sup> | Ethyl acetate | 89.0602 | 0.0005 | X | Oxygen-containing |
| **95.0010** | (C<sub>2</sub>H<sub>6</sub>S<sub>2</sub>)H<sup>+</sup> | Dimethyl disulphide (DMDS) | 94.9989 | 0.0021 | X | Sulphur-containing |
| **109.0760** | (C<sub>6</sub>H<sub>8</sub>N<sub>2</sub>)H<sup>+</sup> | 2,5-dimethylpyrazine | 109.0765 | 0.0005 | X | Nitrogen-containing |
| **118.0651** | (C<sub>6</sub>H<sub>7</sub>N)H<sup>+</sup> | Indole | 118.0656 | 0.0005 | X | Nitrogen-containing |
| **126.9690** | (C<sub>2</sub>H<sub>6</sub>S<sub>3</sub>)H<sup>+</sup> | Dimethyl trisulphide (DMTS) | 126.9710 | 0.0010 | X | Sulphur-containing |
| **137.1325** | (C<sub>10</sub>H<sub>16</sub>)H<sup>+</sup> | Monoterpene | 137.1330 | 0.0005 | X | Hydrocarbon* |

### Effects of salt on the total MVOC emissions

We cultured the bacterial strains under three salt conditions: 0% w/v added NaCl (low salt), 5% w/v (moderate salt), and 10% w/v (high salt). To quantify total net emissions, we summed the mixing ratios (parts-per-billion by volume, ppb_v_) of all signals at each time point (**Fig. 1**).

**Figure 1.**
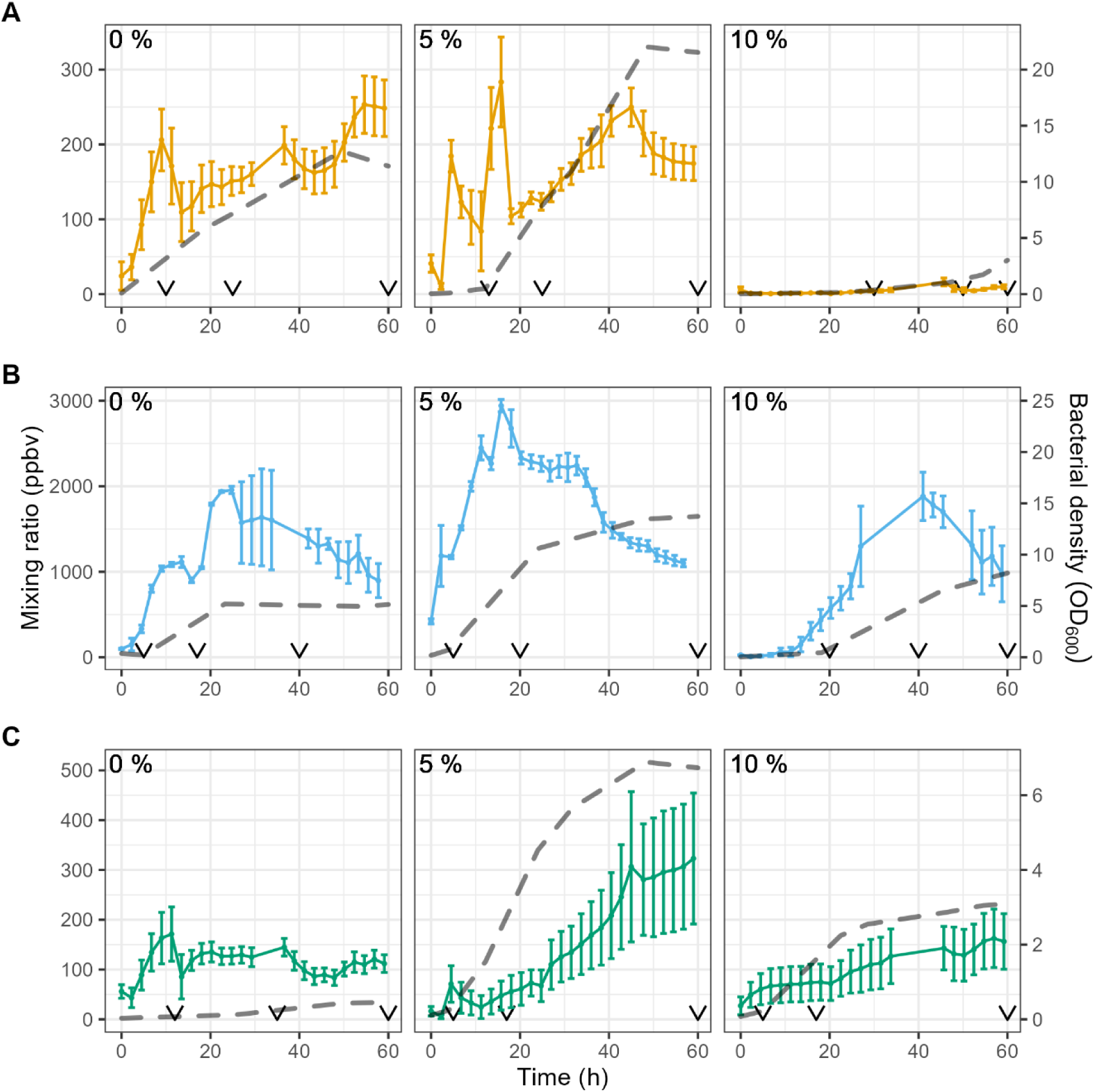
Net headspace total MVOC concentrations and bacterial growth over 60 hours incubation in different salt conditions. The figure shows the strains *Arthrobacter sp.* (**A**, yellow), *N. aurantiaca* (**B**, blue) and *Oceanobacillus sp.* (**C**, green). Datapoints and solid lines: Averaged (n= 3, except fo *N. aurantiaca* in 5% salt, where n = 2) ) summed mixing ratio (ppb_v_) ± standard error. Dark grey dashed lines: Averaged (n=2) growth curves of parallel cultures in units of absorbance at 600nm. V: Timepoints selected for the early, mid and late phases for further analysis. The different NaCl conditions are indicated on each panel in NaCl % (w/v). Note different y-axis scales for each strain.

*Arthrobacter* sp. showed a gradual increase in total MVOC emissions, reaching the highest headspace concentrations near the stationary phase under both low and moderate salt conditions. Under high salt conditions, however, emissions were much lower and significantly delayed. Growth was also delayed, with the exponential phase beginning only near the termination of the experiment, mirroring the delay in emissions.

*N. aurantiaca* produced the highest total MVOC concentrations among the three strains. Emissions generally peaked during the exponential phase, after which they decreased. Under low salt conditions, the highest concentrations occurred closer to the stationary phase than in the other salt conditions.

*Oceanobacillus* sp. showed similar trends in moderate and high salt conditions, with total MVOC concentrations gradually increasing throughout the experiment. On the other hand, total concentrations in low salt were highest early in the experiment, gradually decreasing thereafter.

### Changes in MVOC classes across salt conditions

We compared the relative abundances of different MVOC classes at three time points corresponding to the onset of the exponential phase (“early”), middle of the exponential phase (“mid”), and the stationary phase (“late”), although we note that some cultures did not reach the stationary phase (, see **Fig. 1** for the time points). Notably, each strain showed distinct compositional shifts in response to salt concentration and growth phase (**Fig. 2**).

**Figure 2.**
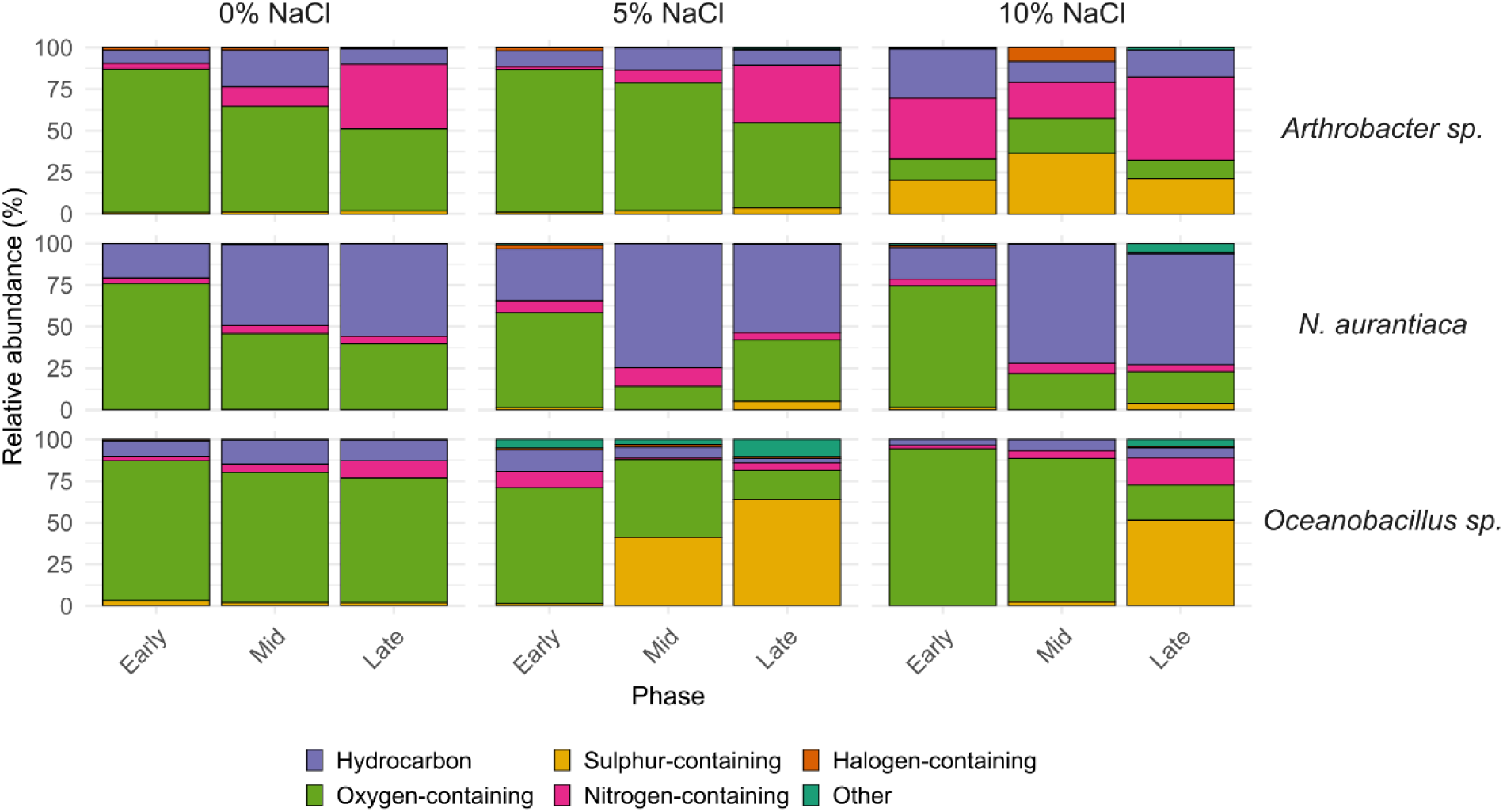
Relative abundances of MVOC classes in the early, mid, and late growth phases. The values shown were calculated after averaging the replicates (n= 2 or 3). The signals were summed into compound classes of MVOCs based on their tentative identities.

Under low salt conditions, all three strains predominantly emitted oxygen-containing compounds in the early phase. *Arthrobacter* sp. and *Oceanobacillus* sp. exhibited similar profiles, with oxygen-containing compounds accounting for over 50% of the total MVOCs, primarily due to acetaldehyde (signal 45 u) and acetone (signal 59 u), respectively. In addition, *Arthrobacter* sp. produced rising concentrations of nitrogen-containing compounds as time passed, mainly 2,5-dimethylpyrazine (signal 109, see also **Fig. 3, panel A3**). In contrast, *N. aurantiaca* showed a marked shift from oxygen-containing compounds (75% of total VOCs) in the early phase to hydrocarbons (55% of total VOCs) in the late phase, with isoprene (signal 69) and an alkenyl fragment (signal 41 u) being the main contributors.

**Figure 3.**
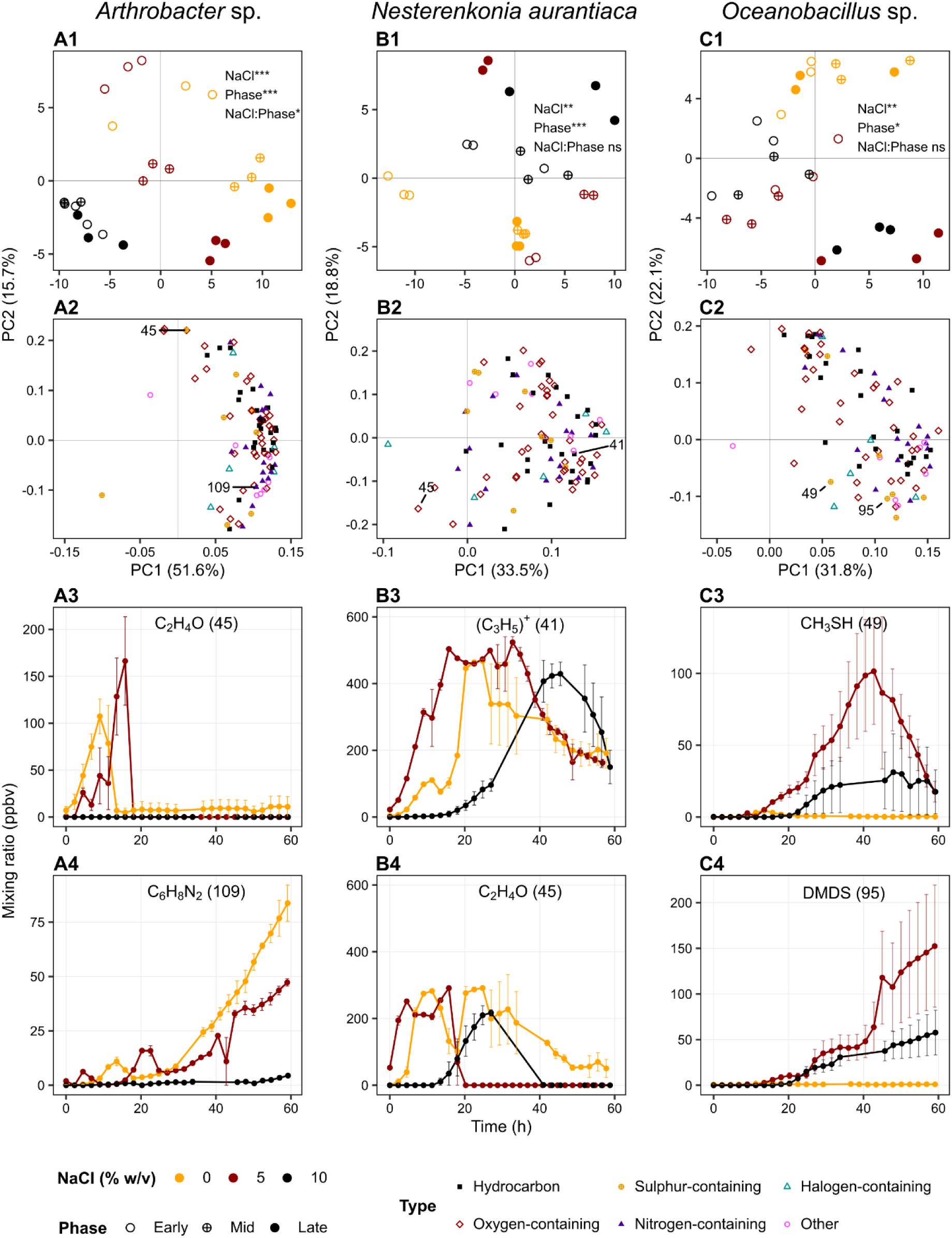
Principal Component Analysis of VOC emission blends for the three bacterial strains across salt treatments and selected phases. The plots correspond to *Arthrobacter sp.* (**A**), *N. aurantiaca* (**B**) and *Oceanobacillus sp*. (**C**). In the PCA score plots (**A1, B1, C1**), the shape of the symbols corresponds to the phase, and the colour corresponds to the NaCl treatment. The explained variances are shown between parentheses for each principal component. The plot annotation shows the significance level of each factor as determined with PERMANOVA (Bray-Curtis dissimilarity) on the log-transformed emission data, with Bonferroni-adjusted p values <0.001 (***), <0.01 (**), <0.05 (*), and >0.05 (ns, not significant). (**A2, B2, C2**): Loadings of the corresponding PCA. The shapes and colours correspond to different classification of the signals. Loading values for each signal are available in Table S1. (**A3, A4, B3, B4, C3, C4**): Net headspace concentration over time of selected signals labelled in the loading plots and their tentative identification. Molecular formulae show the neutral compound, except for signal 41 u. Standard error bars are shown. The colour corresponds to NaCl treatment. DMDS: Dimethyl disulphide.

In moderate salt conditions, differences between strains became more pronounced. *Arthrobacter* sp. maintained a high abundance of oxygen-containing compounds in the early and mid phases, with a gradual increase in nitrogen-containing compounds as in low salt conditions. *Oceanobacillus* sp. displayed an increase in sulphur-containing compounds, with methanethiol (m/z 49) prevalent in the mid phase and dimethyl disulphide (DMDS, m/z 95) in the late phase. *N. aurantiaca* produced the highest proportion of hydrocarbons in the mid phase (52%), with a distinct hydrocarbon profile relative to low salt conditions, dominated by the fragment ions m/z 41, 43, 57, and 71, and isoprene. Moreover, there was a marked reduction in methylbutanal isomers (m/z 87) relative to low salt. *N. aurantiaca* also produced increasing concentrations of methanol (m/z 33) over time, which appeared at lower concentrations under other salt conditions (**Fig. S1**).

In high salt conditions, both *Arthrobacter* sp. and *Oceanobacillus* sp. produced large proportions of nitrogen- and sulphur-containing compounds throughout the growth phases. 2,5-dimethylpyrazine remained dominant for *Arthrobacter* sp., while *Oceanobacillus* produced primarily methanethiol and DMDS. We note that, due to the delayed growth of *Arthrobacter sp.* in high salt, these results are not directly comparable with those under the other salt conditions. *N. aurantiaca* followed a similar pattern to moderate salt, with the same hydrocarbon fragments appearing in comparable proportions.

Overall, distinct strain-specific trends in MVOC class compositions were observed, with *N. aurantiaca* primarily producing hydrocarbons, *Arthrobacter sp.* favouring nitrogen-containing compounds, and *Oceanobacillus sp*. emitting mainly sulphur-containing compounds in the late growth phases.

### Effects of salt on MVOC blends in different growth phases

We conducted a principal component analysis (PCA) for each strain to assess clustering of the samples from the different salt conditions and the early, mid and late time points based on the MVOC data (individual signal values). The PCAs showed clear separations of salt conditions and phases (**Fig. 3**, see also time-lapse animation in Supplementary Material). Permutational Multivariate Analysis of Variance (PERMANOVA) on the log-transformed mixing ratios showed that both salt condition and growth phase had a strong influence on the MVOC blends of the strains (*p* < 0.05).

Among the three strains, *Arthrobacter sp.* (**Fig. 3A**) showed the clearest separation across salinity (separated along PC1) and growth phase (separated along PC2). PERMANOVA showed that salinity and growth phase explained 39% and 30% of the variance, respectively (*p* < 0.01). The PCA loadings suggested that the early phases under low and moderate salt conditions were characterised by ethanol, formic acid (both signal 47 u), and acetaldehyde (45 u), while the later phases were more correlated with nitrogen compounds, such as 2,5-dimethylpyrazine (109 u), as well as monoterpenes (137 u and 81 u). Methanethiol (49 u) was characteristic of the high salt conditions.

*N. aurantiaca* (**Fig. 3B**) showed a slight separation of growth phases along PC2, except for low salt conditions, where the effect was reversed. The mid and late phases in low salt conditions showed little separation along the first two components, although they did separate along the third component (not shown). PERMANOVA showed that both salinity and growth phase were significant (*p* < 0.01) and explained 14% and 34% of the variance, respectively. Some signals (e.g., 45 u, **Fig. 3, panel B4**) showed similar early production in both low and moderate salt conditions, but a far more rapid decline in concentration under moderate salt conditions, which may explain the lack of separation of the mid and late phases in low salt. In most cases (e.g., 41u and 45 u, **Fig. 3, panels B3 and B4**), signals were actively produced during the exponential phase, and their production was delayed under high salt conditions, consistent with the delayed growth. Lastly, isomers of methylbutanal (signal 87 u) were characteristic for *N. aurantiaca* in low salt condition, and the early phase in moderate salt conditions.

*Oceanobacillus sp.* (**Fig. 3C**) showed a separation of the samples in low salt conditions from the rest. The PERMANOVA results showed that salinity accounted for 22% of the variance (*p* < 0.01), while growth phase was marginally significant (*p* = 0.035) and explained 20% of the variance. PC2 correlated with the phases in moderate and high salt conditions, and also separated low salt conditions from the rest. Several compounds were characteristic of *Oceanobacillus sp.* in low salt, such as isoprene (signal 69 u) and signal 89 u (possibly butyric acid, acetoin, or ethyl acetate). The late phases in moderate and high salt conditions were correlated with sulphur compounds, mainly DMDS (95 u), methanethiol (49 u), and hydrogen sulphide (35 u).

A separate PCA with all three strains under moderate salt conditions (**Fig. S2**) showed a clear separation between strains, with *N. aurantiaca* showing the most distinct MVOC profile. In the moderate salt condition, both strain identity (22% of variance), growth phase (45%) and their interaction (22%) each had a strong influence on the MVOC blend in the headspace (*p* < 0.001).

## DISCUSSION

### Effect of salinity on MVOC emissions

In this study, we investigated the influence of salinity on the MVOC production of three bacterial strains isolated from polar desert environments in northern Greenland. Our results show that salinity modulates both the quantity and composition of MVOC production.

While the three strains exhibited distinct volatilomes, we observed some general patterns. *N. aurantiaca*’s total headspace concentrations were highest during the exponential phase in all three salt conditions and decreased in the stationary phase, suggesting that most MVOCs produced by this bacterium are related to central metabolism and growth. In contrast, *Oceanobacillus* sp. and *Arthrobacter* sp. showed highest total concentrations in the stationary phase, suggesting that the MVOC emissions in these bacteria are mainly associated with secondary metabolism. In natural environments, microbial cells are commonly in the stationary phase under nutrient-limited conditions, with sporadic growth periods when nutrients become available^13^. Thus, MVOC production can be highly variable.

We observed a short, sharp peak of acetaldehyde during the exponential phase of *N. aurantiaca* and *Arthrobacter* sp. (**Figs. 3**, panels **A3** and **B4**), as well as a later peak of methanethiol produced by *Oceanobacillus* sp. (**Fig. 3**, panel **C3**). These compounds are commonly emitted by soils during rewetting experiments. For example, Pugliese et al^14^ observed a sharp peak of carbonyl compounds, including acetaldehyde, acetone, butanone and pentanone following tropical soil rewetting after a drought period. The authors attributed the emission of these carbonyl compounds to abiotic reactions based on their short timing. However, the sharp peaks of acetaldehyde that we observed suggests that bacteria can quickly produce similar peaks when they are most metabolically active. Although our setup is not directly comparable to soil conditions (e.g. the three strains were incubated under high initial nutrient availability), this observation suggests that a biotic component may contribute to short-term acetaldehyde emissions following soil rewetting over longer time periods in Northern Greenland.

Under salt conditions that severely impaired growth (i.e., *Arthrobacter* sp. in high salt, and *Oceanobacillus* sp. in low salt), the volatilomes were most distinct compared to other salt conditions. Furthermore, net headspace concentrations only started to increase toward the end of the experiment for *Arthrobacter* sp. under high salt, limiting comparability with other treatments. This suggests that inhibitory salinity concentrations not only suppress overall metabolic output but may also lead to shifts in biochemical pathway usage consistent with stress responses which influence MVOC production. In the case of *N. aurantiaca*, we observed a different mixture of hydrocarbons and methylbutanal isomers in moderate and high salt compared to low salt, suggesting metabolic changes. Salt stress has a profound effect on metabolism, upregulating – among others – genes related to ion transport and osmoprotectant production^15^. In the halotolerant bacterium *Staphylococcus aureus*, growth in very high salt conditions (20% w/v) upregulates genes related to sulphur metabolism^16^, which may lead to altered production of sulphur-containing MVOCs, as observed in our study for *Oceanobacillus* sp. in moderate and high salt conditions.

Overall, our results suggest that both the total MVOC quantities produced, and their chemical makeup may serve as indicators of microbial stress and salt tolerance, reflecting strain-specific physiological adaptations to saline environments. Thus, our results agree with hypotheses (1) and (2).

### Salinity-Driven Changes in Bacterial Metabolism

The observed changes in MVOC composition under different salt conditions likely reflect underlying shifts in metabolic activity. In this section, we examine how salinity influenced the concentrations of specific MVOCs, with a focus on compounds linked to stress responses and secondary metabolism.

*Arthrobacter sp.* produced significant quantities of 2,5-dimethylpyrazine, particularly under low and moderate salt conditions in the late growth phase. *Oceanobacillus sp.* also produced this compound across all salt conditions in the late phase. Pyrazines can be produced by Maillard reactions between carbohydrates and amino acids at high temperatures; however, they are ubiquitous in nature^17^. Pyrazines, including 2,5-dimethylpyrazine, exhibit antimicrobial activity against phytopathogenic bacteria^18^. In the case of *Arthrobacter sp.*, this pyrazine may provide a competitive advantage in its native permafrost habitat. The concentration of 2,5-dimethylpyrazine was nearly double in low salt compared with moderate salt despite lower growth in low salt, suggesting that this compound may be part of a broader metabolic response to salt stress, with protective or competitive functions. Previous studies have shown that salinity changes influence the secondary metabolism of marine bacteria^19^. The large production of pyrazines is surprising since terrestrial Arctic ecosystems in general are considered nitrogen limited^20^ and pyrazine emissions may lead to ecosystem nitrogen loss. However, pyrazine emissions from bacteria may also lead to nitrogen re-location, e.g., from thawed permafrost soil to surface soils where pyrazines and other nitrogen-containing MVOCs may be taken up by microorganisms.

The hydrocarbon fragments produced by *N. aurantiaca* may partly originate from 1-butanol (signal 57 u), as well as 2- and 3-methylbutanol (43 u and 71 u)^12^, all of which were detected via GC-MS. Both 2- and 3-methylbutanol are commonly produced by actinobacteria^21^ and are likewise emitted by bacteria associated with desert plants^22^. Other strains in the genus *Nesterenkonia* have been shown to produce butanol via the acetone-butanol-ethanol fermentation pathway^23^, although this pathway is unlikely to be the origin of the observed butanol because the bacteria were grown under aerobic conditions. Furthermore, we only detected small quantities of ethanol (47 u) in the headspace. 2- and 3-methylbutanol can also be produced from isoleucine and leucine, respectively, by deamination and decarboxylation that yield 2- and 3-methylbutanal, which are then reduced by a NAD(P)-dependent alcohol dehydrogenase to form alcohols^24^. The composition of the medium, rich in amino acids, may have favoured this pathway. The net concentration of methylbutanal isomers remained high under low salt conditions but decreased rapidly under moderate and high salt conditions. As the methylbutanal signal decreased, methylbutanol increased, suggesting that methylbutanal reduction may be inhibited in low salt conditions in this halophilic bacterium. In the cyanobacterium *Synechocystis spp.*, salt and osmotic stress increase the expression of an alcohol dehydrogenase that preferentially reduces aldehydes to alcohols^25^. Furthermore, alcohol dehydrogenases are implicated in cold and salt stress responses in plants^26^.

Notably, only *N. aurantiaca* produced methanol, primarily in moderate salt conditions, where its concentration increased steadily throughout the experiment. Methanol is a ubiquitous MVOC that is both emitted and consumed by soil microorganisms^27^. In the Arctic region, methanol is emitted from thawing permafrost^4^. Major microbial sources of methanol include the breakdown of pectin from dead plant material in soil^28^ and the oxidation of methane by methanotrophs, where methane is oxidised sequentially to methanol, formaldehyde, formic acid and finally CO_2_^29^. However, these processes could not have been the source of methanol in our experiment, as the growth medium did not contain pectin, and the zero-air used as the headspace gas did not contain methane. Methanol can also be produced by the hydrolysis of methyl acetate^30^, although we did not detect significant amounts of either methyl acetate or acetic acid.

*Oceanobacillus sp.* produced significant amounts of methanethiol and DMDS under moderate and high salt conditions. As this strain grows optimally at approximately 7.5% w/v NaCl^31^, its diminished growth and lack of sulphur compound production under low salt conditions suggest that the biochemical pathways involved may be inhibited by osmotic stress. Methanethiol is typically derived from sulphur-containing amino acids methionine and cysteine or from demethylation of the osmoprotectant dimethylsulfoniopropionate^32^. DMDS arises either via abiotic oxidation of methanethiol^33^ or via chemical disproportionation of S-methyl methanethiosulfinate, a metabolite found in plants^34^ and marine bacteria^35^. Both methanethiol and DMDS are widespread in terrestrial and marine environments^32^ and play various ecological roles; for instance, DMDS, exhibits antifungal activity^36^. In estuarine sediments, Magalhães et al.^5^ found that salinity correlates with methanethiol formation^5^. As the authors observed formation of methanethiol mainly under anoxic conditions, they hypothesised that oxidation to DMDS under oxic conditions prevented methanethiol detection. While methanethiol is a major component of the Arctic Ocean sulphur budget^37^, its emission from thawing permafrost or polar deserts has not been reported. Methanethiol oxidation in the atmosphere eventually leads to the formation of sulphuric acid droplets and cloud condensation nuclei, with potential environmental implications^38^.

Salinity influences the solubility and partitioning of volatile compounds between liquid and gas phases, a phenomenon known as the “salting-out effect”, which may have enhanced the detection of some MVOCs in the headspace as salt concentration increased. However, the salting-out effect alone is insufficient to explain the trends observed. In some cases, higher concentrations were observed under low compared to moderate salt conditions, such as 2,5-dimethyl pyrazine in *Arthrobacter sp.* Given that alcohols are especially affected by the salting-out effect^39^, the production of 2- and 3-methylbutanol by *N. aurantiaca* was likely underestimated under low salt conditions. Nonetheless, the methylbutanal decline over time in moderate and high salt conditions strongly indicates active metabolism of methylbutanal, which was absent under low salt conditions. Dehydrogenase-dependent reduction to alcohols is a likely pathway, although other metabolic routes may have contributed to the loss of methylbutanal isomers.

Our findings represent a first characterization of the effect of salinity on the production of MVOCs in bacteria from Arctic desert soils, offering insights into how osmotic stress affects microbial volatile emissions. The increased production of sulphur-containing volatiles by *Oceanobacillus sp.* under high salt conditions, along with the variability in methylbutanal and methylbutanol emissions in *N. aurantiaca*, highlights how this stress influences microbial metabolism. These results also raise key questions for future research. We observed a significant production of sulphur- and nitrogen-containing MVOCs, yet their emission in these remote areas remains uncharacterised. With the Arctic warming two to three times faster than the global average, increased extreme weather events are likely to cause soil salinity fluctuations^10^. Consequently, microbial activity in geochemical cycles may increase, altering MVOC emissions potentially leading to enhanced ecosystem nutrient losses through element mobilization. These findings highlight the potential of MVOC profiling as a tool for assessing microbial stress responses in Arctic environments and providing a foundation for exploring the role of volatiles in cold-region nutrient cycling.

## MATERIALS AND METHODS

### Bacterial strains and growth conditions

The strains used in this study were isolated and characterised in a previous study^31^. Briefly, the strains were isolated using saline media from samples collected in Peary Land, Northern Greenland, north of 82°N: biological soil crust from Citronen Fjord (*Oceanobacillus sp.* CF4.6), biological soil crust from Cape Morris Jessup (*N. aurantiaca* CMS1.6), and permafrost soil from the Kap København formation (*Arthrobacter sp.* KK5.5). Based on the response of the strains to NaCl (**Fig. S3**), we decided to use three different NaCl concentrations: 0, 5 and 10% w/v (low, moderate and high salt, respectively). The medium used for all three strains was a modified HM medium^40^ containing (g L^-1^): NaCl (0, 50 or 100); MgSO_4_·7H_2_O (1); CaCl_2_·2H_2_O (0.36); KCl (2); KBr (0.27); FeCl_3_·6H_2_O (0.002); NaHCO_3_ (0.06); Proteose-peptone no. 3 (5); Yeast Extract (10); and glucose (1), adjusted to pH 7,8-8. For solid media, Bacto agar (20 g L^-1^) was added.

In all cases, the strains were grown at 25 °C and constant agitation at 180 rpm. Liquid cultures (20 mL) were grown in 350-mL jars. The cultures were inoculated with 600 µL of overnight cultures with the same NaCl concentration; these precultures were prepared by inoculating 5 mL of medium with a colony picked after 24-48 h of growth on agar. Uninoculated HM medium at each NaCl concentration was used as negative controls.

To avoid disturbing the online MVOC sampling, two additional replicates were grown in parallel under identical conditions to allow growth monitoring via OD₆₀₀ measurements at regular intervals. To avoid saturation, samples were diluted in fresh medium when optical density was above 0.5 units.

### MVOC sampling and analysis

#### Offline and online sampling

Offline sampling of volatiles from the headspace of the bacterial cultures incubated at 25 °C and negative control samples was done *via* sorbent tubes (C2-AXXX-5032 Hydrophobic, Tenax TA/Carbograph 1TD, Markes International, UK) with a flow rate of 200 mL min^-1^ and a sampling time of 5-10 min (**Fig. 4A**). The sorbent tubes were stored at 4 °C until analysis. Headspace sampling was done during exponential growth, ranging between 22-54 h depending on the strain and salt concentration.

**Fig 4.**
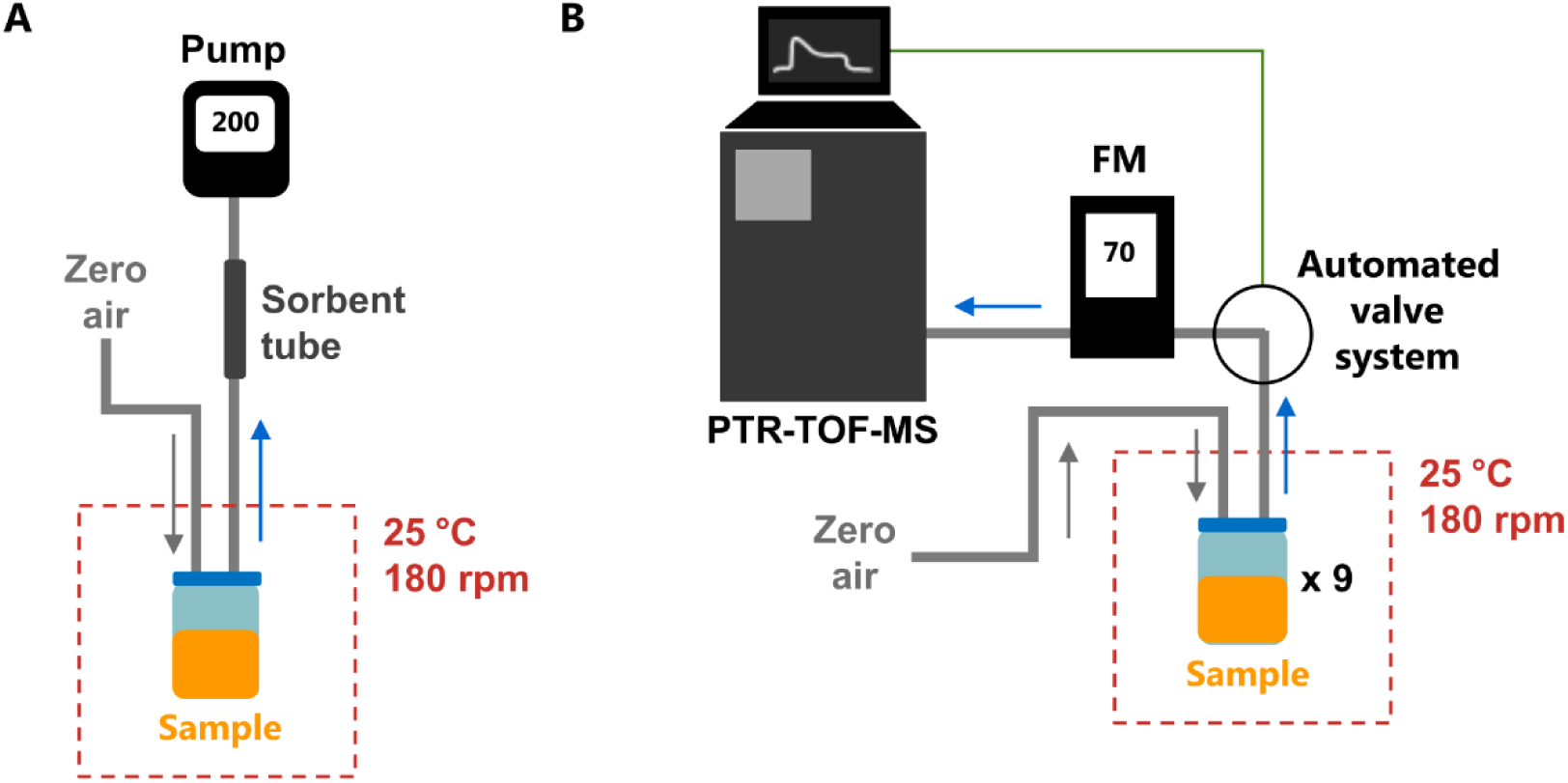
Schematic representation of the (A) offline, and (B) online sampling setups. Grey lines indicate PTFE tubing, and green line digital connection. A handheld pump (Pocket Pump TOUCH, SKC, PA, US) was used for pumping the sample headspace through the sorbent tube at a flow rate of 200 mL min^-1^. A mass flow meter (FM, Alicat Scientific, AZ, US) was used to ensure sufficient flow through the online system and into the measurement instrument.

Online sampling of volatiles from the bacterial culture headspace was performed using an automated solenoid valve system with a polytetrafluoroethylene (PTFE) sampling line connected to the PTR-TOF-MS instrument (**Fig. 4B**). Each sample was connected to the sampling line with a flow rate of 70 mL min^-1^. Each sample was successively measured for 15 min until concentrations stabilized, and the last two minutes of the sampling were used for data analysis. An automated solenoid valve system rotated between the samples throughout the 60 h culture period.

Four replicates were used for offline sampling and three for online sampling.

#### Gas chromatography**–**mass spectrometry (GC**–**MS)

The analysis of the MVOCs captured by the sorbent tubes was performed after thermal desorption by GC-MS as described previously^41^. PARADISe (Chemometrics Research)^42^ was used for the chromatogram and mass spectral analysis with the National Institute of Standards and Technology mass spectral library (NIST14)^43^. Standards were measured alongside the samples for the quantification and identification of the 45 individual compounds listed in **Table S2,** with all other identities considered tentative.

#### Proton-transfer-reaction time-of-flight mass spectrometry (PTR-ToF-MS)

A PTR-ToF-MS instrument (PTR-TOF 1000 *ultra*, Ionicon, Austria) with a specified mass resolution of >1500 was used for the continuous, online VOC measurements. Hydronium (H_3_O^+^) was used as the reagent ion. Measurements were performed between m*/z* 17–239, with one spectrum measured every 5 seconds.

The operating conditions were as follows: drift tube voltage of 500 V (corresponding to reduced electric field *(E/N*) of 116 Td); drift tube pressure of 2.20 mbar; drift tube temperature of 60 °C; H_2_O flow of 6 standard cubic centimetres per minute (sccm); ion source current of 3 mA; inlet flow of 50 sccm; inlet temperatures of 70 °C. The PTR-ToF-MS measurement software (ioniTOF v4.0; Ionicon) was used for monitoring measurements in real-time, as well as for the automated control of the solenoid valves rotating between different sample headspaces. The PTR-ToF-MS operation conditions were chosen as to minimize the formation of unwanted water clusters in high humidity samples and simultaneously prevent excess fragmentation of sample molecules^44^.

PTR-MS Viewer v3.4.4.13 (Ionicon, Austria) was used for spectral analysis and conversion of raw measurement data to mixing ratios (ppb_v_). The PTR-ToF-MS instrument uses an internal standard (1,3-diiodobenzene) and the H_3_^16^O^+^ signal for two-point mass scale calibration. A standard gas mixture (Air Liquide, Denmark) was used to calibrate the concentrations of ten VOCs (**Table S3**) and for transmission curve measurements. We note that PTR-ToF-MS can distinguish between some isobars (same nominal mass), but not isomers (same exact mass). We tentatively identified compounds by comparing the measured mass of a signal to the theoretical mass of the assumed molecular formula, with the tentative identity deemed acceptable if the mass difference was < 0.01 u. We also used the isotopic analysis tool in PTR-MS Viewer v3.4.4.13 to further confirm signal identities, compounds of interest were further identified by comparing the PTR-ToF-MS data to the GC-MS data.

### Data preparation and statistical analysis

Unless otherwise specified, data processing and statistical analysis were performed using R version 4.3.3. To prepare the GC-MS data for analysis, the negative controls were evaluated for outliers, averaged, and subtracted from the samples separately for each salt treatment. For the PTR data, the negative controls were evaluated for outliers, averaged, and then smoothed using locally estimated scatterplot smoothing (LOESS) with a span of 0.3 to remove before subtracting them from the samples for each salt treatment and sampling run. Negative values in both datasets were set to zero.

Three time points (early, mid, and late phases) were selected based on growth curves from parallel cultures (**Fig. 1**). Concentrations at these three time points were then used in SIMCA 17 (Umetrics, Umeå, Sweden) to perform Principal Component Analysis (PCA). Prior to the PCA, the data was log-transformed using the Auto-transform feature.

Permutational multivariate analysis of variance (PERMANOVA)^45^ was performed on the log-transformed concentrations at the three selected phases using the *adonis2* function in the R package *vegan*^46^ version 2.6-6.1. The factors were Strain, NaCl, and Phase. Bray-Curtis dissimilarity was used for the analysis, with 9999 permutations. Replicates were treated as strata to account for repeated measurements of the same samples, ensuring permutations respected the experimental design. Bray-Curtis dissimilarity was chosen for its sensitivity to relative differences and particular suitability for datasets containing many zeroes.

## Supporting information

Supplement and description of suplpementary files

Supplementary animation

## ACKNOWLEDGEMENTS

We thank Laura Meredith for many fruitful discussions about the analysis of the data. This project was funded by the Novo Nordisk Foundation under the NNF Interdisciplinary Synergy Program (grant number. NNF19OC0057374): Effects of bacteria on atmospheres of Earth, Mars, and exoplanets -- adapting and identifying life in extraterrestrial environments. This project was also supported by Jenny and Antti Wihuri Foundation (grand number 220323), and The Danish National Research Foundation within the Center for Volatile Interactions (VOLT, DNRF168). The bacteria in this study were isolated from samples collected during the Leister Around North Greenland 2021 Expedition financed by the Leister Stiftung, Switzerland. We thank the Greenlandic authorities for granting permission to sample in the Northeast Greenland National Park (Expedition Permit C-21-549) and to export the soil samples to Denmark for scientific work (Non-exclusive license number G21-031 for utilization of Greenlandic genetic resources).

## AUTHOR CONTRIBUTIONS

M.A.S.G. and A.P. conceptualised this study. M.A.S.G. grew the strains and collected GC-MS data. K.R. prepared and optimised the PTR-MS setup. K.R., M.A.S.G., and M.B.R. performed the PTR-MS measurements and data collection. K.R processed the raw PTR data. M.A.S.G. and K.R. analysed the VOC data. M.A.S.G performed statistical analyses. A.P. and R.R. provided supervision. M.A.S.G. wrote the original draft and produced the figures. K.R drafted the MVOC sampling and analysis section and related figures. All authors revised and contributed to the drafted manuscript.

## DATA AVAILABILITY

The data collected and used in this study is available at [link will be here in final version].

## CONFLICTS OF INTEREST

The authors have no conflicts of interest to declare.

## Notes

### Competing Interest Statement

The authors have declared no competing interest.

