## Supplement and description of suplpementary files for "Volatilomic complexity of three Northern Greenland bacterial isolates across a salt gradient"

This file includes supplementary figures, small tables, and description of supplementary animation. Tables S1 and S4 are available as a separate .xlsx file. Supplementary animation is available as a separate .GIF file.

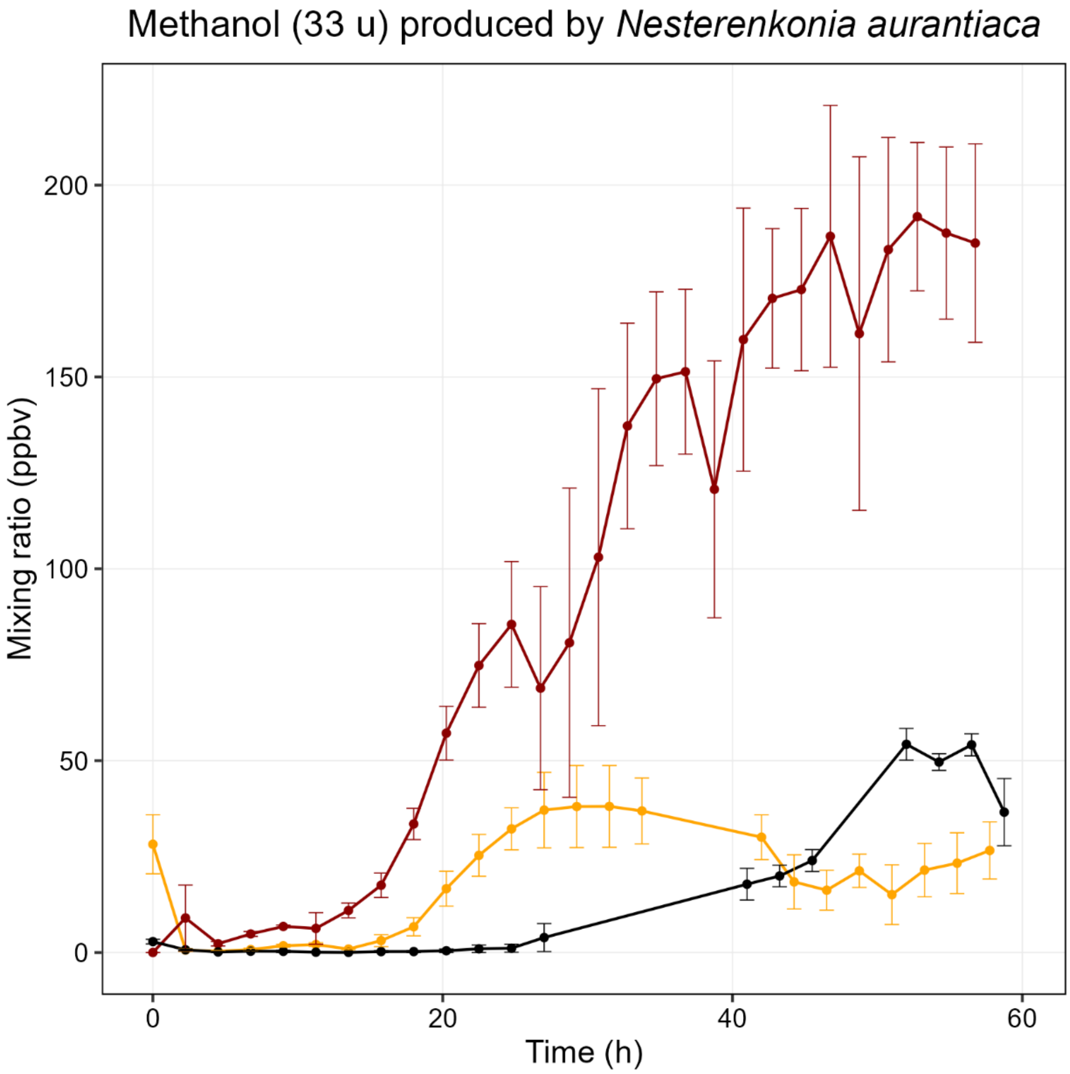


**Figure S1. Methanol produced by *Nesterenkonia aurantiaca*.** Colours indicate low salt (yellow), medium salt (red) and high salt (black).

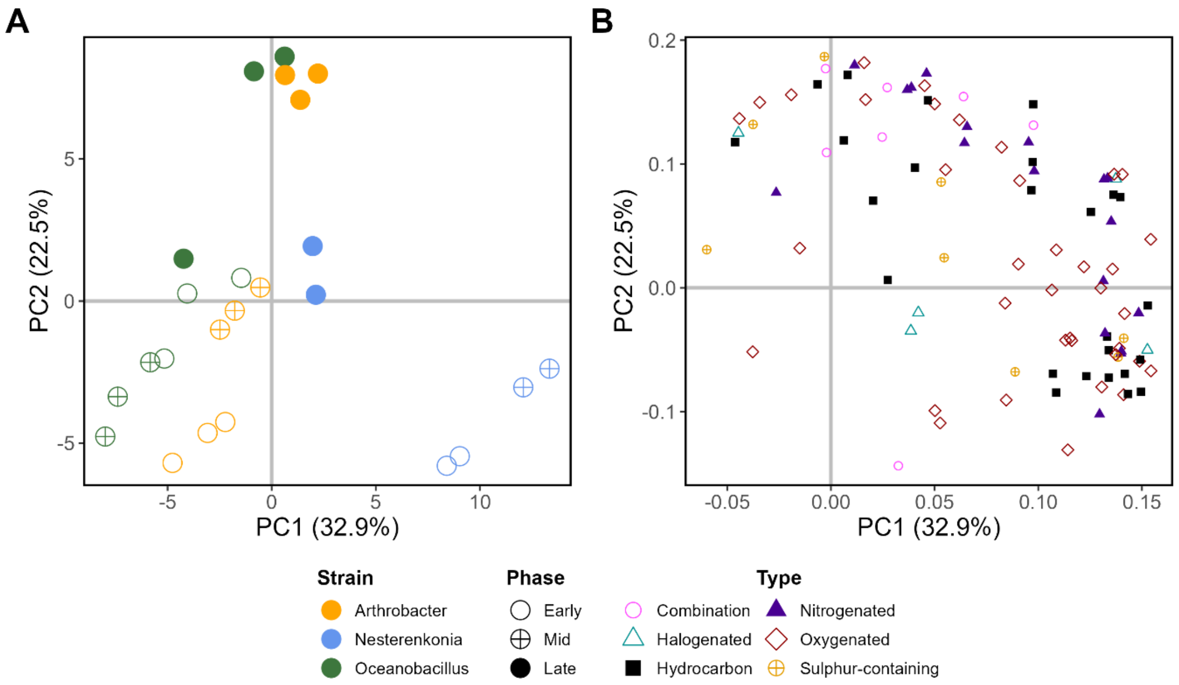


**Figure S2. PCA of the MVOCs produced by the strains in moderate (5% w/v) NaCl conditions detected using PTR-ToF-MS.** Plot A shows the scores and plot B, the loadings of signals. The explained variances are shown between parentheses for each principal component. Loading scores are available in Table S2.

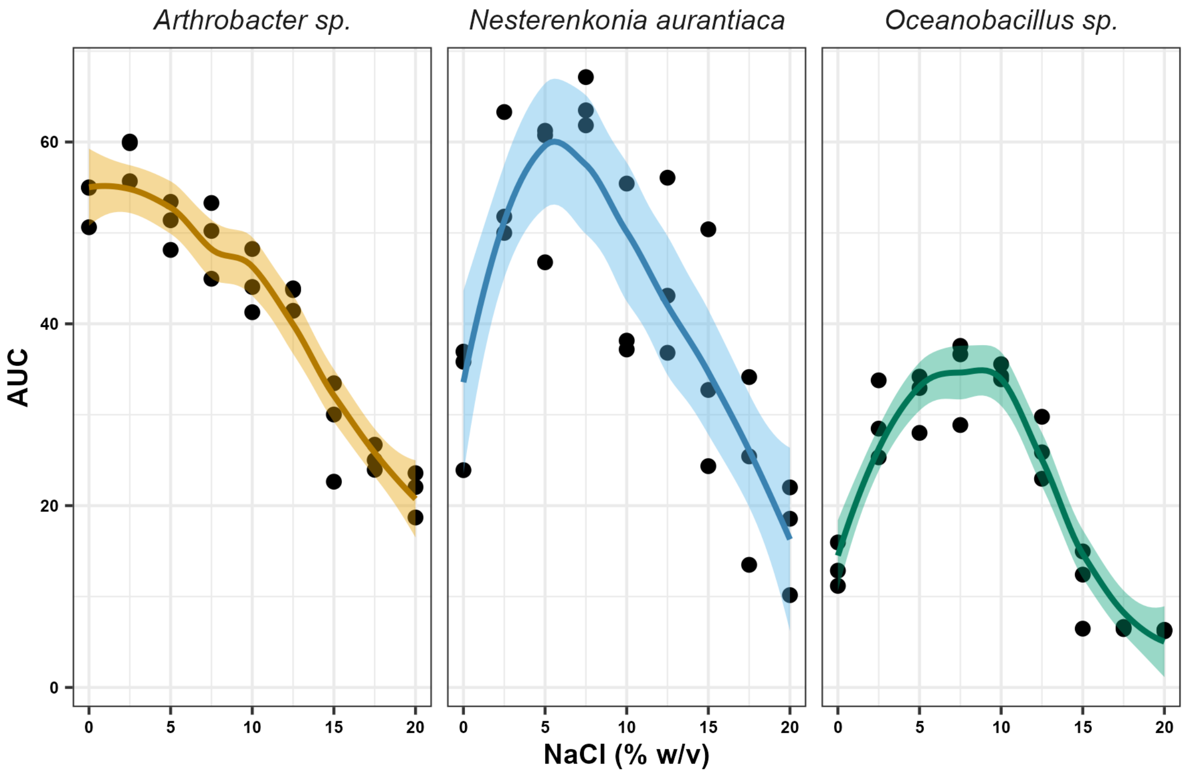


**Figure S3. Effect of NaCl on the growth of the strains used in this study.** AUC: Area under the curve at time = 60 hours, calculated using the trapezoid rule after fitting optical density data to a Gompertz model. Dots: Individual samples. Solid lines: LOESS fit lines calculated with span = 0.5. Shaded area: 95% confidence interval. Data and methodology from additional ref (1).

**Table S1 (separate excel file).** Loading scores of the Principal Component Analyses presented in Fig. 3 of the main text. In addition, loading scores of the PCA presented in figure S2 are also presented. Dashes are signals that were not included in the PCA due to lack of variability between samples (typically because all values were 0), as recommended by the software used.

**Table S2.** Compounds used in TD-GC-MS standard measurements.

| **GC-MS, Standard compounds** | | |
|---|---|---|
| isoprene | cis-3-hexenyl acetate | (plus)-a-longipinene |
| 2-butanone | a-phellandrene | (minus)-isolongifolene |
| hexane | 3-carene | b-caryophyllene |
| 2-methylfuran | p-cymene | a-humulene |
| ethyl acetate | d-limonene | caryophyllene oxide |
| toluene | benzyl alcohol | 3-methyl-1-butanol |
| 1-octene | eucalyptol | 2-methyl-1-butanol |
| hexanal | ocimene | |
| 2-furaldehyde | y-terpinene | |
| trans-2-hexen-1-al | acetophenone | |
| cis-3-hexen-1-ol | terpinolene | |
| p-xylene | linalool | |
| o-xylene | nonanal | |
| a-pinene | (minus)-a-thujone | |
| camphene | (plus)-camphor | |
| benzaldehyde | (minus)-borneol | |
| 1-octen-3-ol | cis-3-hexenyl butyrate | |
| b-pinene | methyl salicylate | |
| myrcene | bornyl acetate | |
| octanal | indole | |

**Table S3**. Compounds used in PTR-TOF-MS standard measurements.

| **PTR-TOF-MS, Standard compounds** | |
|---|---|
| Methanol | Butanone |
| Acetonitrile | Benzene |
| Acetaldehyde | Toluene |
| Acetone | m-Xylene |
| Isoprene | a-Pinene |

**Table S4 (separate excel file). Loading scores of the Principal Component Analyses presented in Supplementary Animation.** Dashes are signals that were not included in the PCA due to lack of variability between samples (typically because all values were 0), as recommended by the software used.

**Supplementary animation (separate .GIF file)**. This PCA animation shows the progression of the cultures over 60 hours along PC1 and PC2. To produce this animation, a separate PCA for each strain was performed using all time points as described in the main text. The data points were smoothed using LOESS (step = 0.3) to avoid excessive jittering. Loading scores are available in Table S4.
