## Supplementary figures and images for "Volatilomic complexity of three Northern Greenland bacterial isolates across a salt gradient"

### Supplementary animation

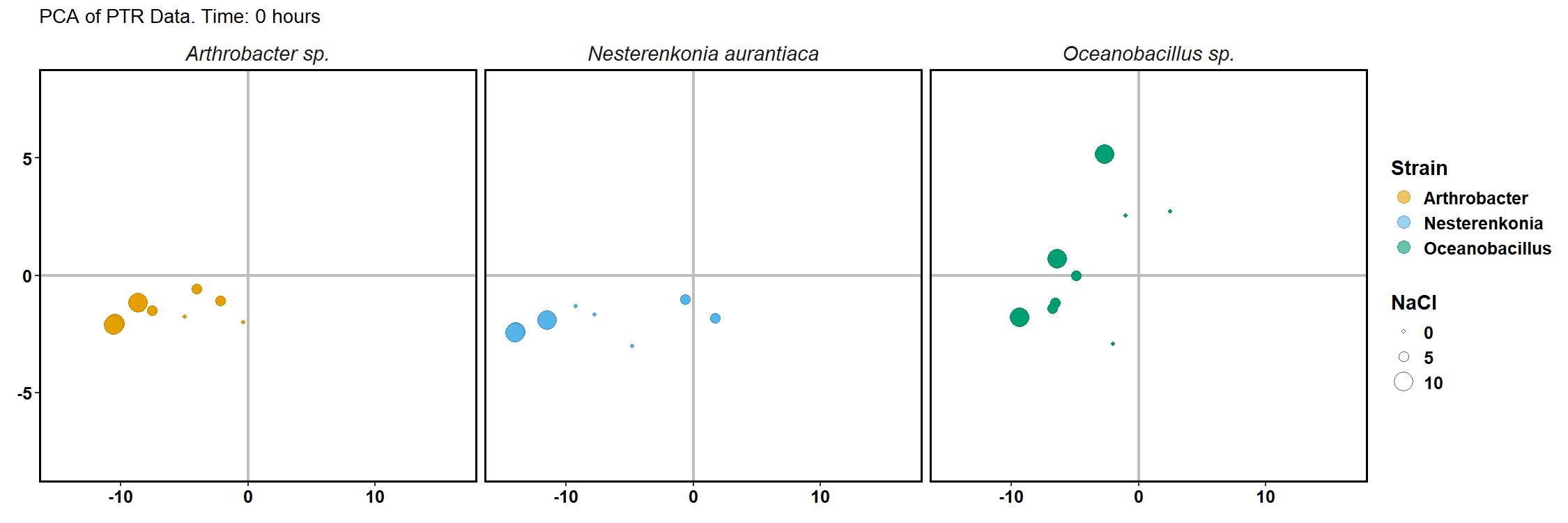
